# The CD4+ T cell regulatory network mediates inflammatory responses during acute hyperinsulinemia: a simulation study

**DOI:** 10.1101/058743

**Authors:** M.E. Martinez-Sanchez, M. Hiriart, E. R. Alvarez-Buylla

**Affiliations:** Departamento de Ecología Funcional, Instituto de Ecología, Universidad Nacional Autónoma de México.; Centro de Ciencias de la Complejidad, Universidad Nacional Autónoma de México.; Departamento de Neurodesarrollo y Fisiología, Instituto de Fisiología Celular, Universidad Nacional Autónoma de México.

## Abstract

Obesity is linked to insulin resistance, high insulin levels, chronic inflammation, and alterations in the behavior of CD4+ T cells. Despite the biomedical importance of this condition, the system-level mechanisms that alter CD4+ T cell differentiation and plasticity are not well understood. We model how hyperinsulinemia alters the dynamics of the CD4+ T regulatory network, and this, in turn, modulates cell differentiation and plasticity. Different polarizing micro-environments are simulated under basal and high levels of insulin to assess impacts on cell-fate attainment and robustness in response to transient perturbations. In the presence of high levels of insulin Th1 and Th17 become more stable to transient perturbations and their basin sizes are augmented, IL10 producing regulatory T cells become less stable or disappear, while TGFB producing cells remain unaltered. Hence, the model provides a dynamic system-level explanation for the documented apparently paradoxical role of TGFB in both inflammation and regulation of immune responses and the emergence of the adipose Treg phenotype. Furthermore, our simulations provide novel predictions on the impact of the micro-environment in the coexistence of the different cell types, proposing that in pro-Th1, pro-Th2 and pro-Th17 environments effector and regulatory cells can coexist, but that high levels of insulin severely affect regulatory cells, specially in a pro-Th17 environment. This work provides a system-level formal and dynamic framework to integrate further experimental data in the study of complex inflammatory diseases.

## Introduction

Obesity-associated chronic inflammation is a complex phenomenon that results from the interaction between adipose tissue, hyperinsulinemia, and chronic inflammation (1–4). Together, these linked conditions increase the risk to develop metabolic syndrome and type 2 diabetes mellitus. To understand how such complex syndrome emerges, it is necessary to use an integrative, system-level and dynamic approach that takes into consideration: the non-linearity of the interactions, the strong effect of the environment, the constant crosstalk and feed forward interactions among the genetic and non-genetic components involved, and the synchronic or concerted nature of various regulatory events and conditions involved occur (5–8). Most studies have focused on the direct relationship between macrophages and obesity (9), meanwhile, important questions concerning the relationship between obesity, insulin, and CD4+ T cell types populations and plastic changes among them remain unaddressed. These probably play important roles in the onset of inflammatory responses, and their systemic impact remains unresolved. A starting point, involves understanding: (i) the complex regulatory network involved in the cell fate attainment of CD4+ T cell types (10–12), (ii) how such network responds to extracellular metabolic and environmental conditions (13–14) (iii) how the resulting system modulates the inflammatory and immune responses (1–4).

Obesity-associated chronic inflammation result from prolonged excessive nutrient intake (15, 16). Under such condition, adipocytes in the visceral adipose tissue (VAT) stimulate the inflammatory response by producing pro-inflammatory cytokines, increasing activated macrophages, but also altering the CD4+ T cell population which likely feedbacks to inflammation (15,17). This inflammatory response causes a decrease in glucose intake, which affects glucose metabolism and may indirectly promote an increase in insulin production by beta cells (1–4, 15, 16, 18). Hyperinsulinemia is strongly associated with metabolic syndrome that is typified by obesity, hypertension, dyslipidemia, renal failure, fatty liver disease, certain cancers and cardiovascular diseases, among others. Despite the fact that such syndrome is clearly characterized, we still do not understand the system-level underlying mechanisms, as well as the global health consequences associated to hyperinsulinemia (18).

CD4+ T cells are fundamental modulators of immune challenges and the homeostasis of the immune system. Naive CD4+ T cells (Th0) are activated when they recognize an antigen in a secondary lymphoid organ. CD4+ T cells may attain different cell fates depending on the cytokine milieu and other signals in their micro-environment. The cytokines can be produced by the lymphocyte (intrinsic) or by other immune cells (extrinsic). The different cell types express characteristic transcription factors and cytokines and have been associated with specific roles in the immune system (19). The classification of CD4+ T cells in subsets has been complicated, as they are highly heterogeneous and plastic. There are reports of hybrid cells that express transcription factors and cytokines from more than one cell type (20,21), for example. Furthermore, CD4+ T cells can plastically alter their expression patterns in response to environmental conditions (22–24). Such complex and dynamic plastic behavior has started to be explained at the system level using multistable network models (10–12).

Regulatory T cells maintain immune tolerance; regulate lymphocyte homeostasis, activation, and function. Regulatory T cells can be classified into various types. Treg cells are characterized by the transcriptional factor Foxp3, high expression of CD25+, and they produce TGFβ and IL10. But, these two cytokines can also be expressed independently of Foxp3. TGFβ is necessary for the differentiation of regulatory Tregs and effector Th17 cells. TGFβ has a context-specific role in the immune response; it can suppress or enhance the immune reaction, depending on its cofactors (25–27). IL-10 is an immunosuppressive cytokine produced by many cells of the immune response. It acts as a feedback regulator of the immune response by inhibiting the production of inflammatory cytokines (28). Moreover, T cells that express Tbet or GATA-3, in addition to certain regulatory factors, are important in regulating the Th1 and Th2 response (29–31).

CD4+ T cells are involved in the inflammatory feedback loop in obesity-associated tissue inflammation. In the obese VAT murine models and humans, an enrichment of the Th1 and Th17 populations and a decrease in regulatory T cells has been described (3, 32–34). Th1 and Th17 cells produce proinflammatory cytokines that inhibit insulin signaling. The transcriptional profiles and functions of Tregs are also altered, they express proinflammatory cytokines like IFNγ, and IL-10. This change in expression patterns causes Tregs to cluster with inflammatory T cells (3, 32–34). While TGFβ is detectable in adipose tissue, its role in regulating Treg cells is unclear (34). Paradoxically, it has been reported that adipose tissue Treg cells decrease seems to both, improve and worsen insulin resistance (33,35,36). Such behavior could be linked to a multistable dynamic underlying system such as that recently proposed to study CD4+ T cell differentiation and plasticity (10–12). On the other hand, the general metabolic state of an individual also affects CD4+ T cells. Obesity is associated with increased insulin levels, which affects CD4+ T cells. Insulin is necessary for the survival and proliferation of activated CD4+ T cells. Effector T cells, such as Th1, Th2 and Th17, depend on glycolysis, while resting (not activated) regulatory and memory T cells depend mainly on lipid oxidation. But in obese VAT, the high levels of insulin over-activate the AKT pathway, inhibiting IL-10 production and its regulatory functions in CD4+ T cells (14). Hence, the relationship between insulin resistance and CD4+ T cells is still unclear.

We propose here a theoretical simulation study, to explore the molecular interactions between the previously published CD4+ T cell regulatory network (12) and insulinemia (14). Such a system-level mechanistic approach is fundamental for understanding CD4+ T cell differentiation and plasticity dynamics at the cellular level in response to the metabolic state of hyperinsulinemia. We also analyze such altered dynamics of CD4+ T cell dynamics under different IL10 environments. We used the Boolean regulatory network for studying CD4+ T cell differentiation and plasticity dynamics in response to insulin. The system includes transcription factors, signaling pathways, intrinsic and extrinsic cytokines (12), as well as the impact of basal and high levels of insulin (14). The model recovers the differentiation of T CD4+ cells, including effector (Th1, Th2, Th17) and regulatory (iTreg, Th1R, Th2R, Foxp3-IL10+ and Foxp3-TGFβ+ cells) cell types (12,19). Here, we show how hyperinsulinemia shapes CD4+ T cell attainment by reducing the production of IL-10 and causing a shift towards pro-inflammatory, resting, or TGFβ+ producing cell types. Constant pro-regulatory signals can counteract this change. We also explore how the presence of high levels of insulin in the environment alters the plasticity of CD4+ T cell in response to transient fluctuations in the elements of the network. High insulin also favors transitions towards inflammatory, resting or TGFβ+ producing cell types and reduces the stability of regulatory cell types. In this way, we show how the CD4+ T cell molecular network model proposed before (12) seems to mediate the observed cellular behavior in obesity-associated chronic inflammation. This network model constitutes a useful framework to further explore the system-level mechanisms involved in inflammatory conditions including obesity.

## Results

### CD4+ T cell regulatory network

We expanded the previously published T CD4+ cell transcriptional-signaling regulatory network (12) to include the effect of insulin in the differentiation of CD4+ T cells, according to experimental data (14). The CD4+ T cell differentiation/plasticity network focuses in activated CD4+ T cells in VAT, and was grounded on experimental data [File S1]. Using this model we studied the role of the different network components in the cellular dynamics and the impact of the environment in cell fate attainment and plasticity patterns [Figure 1]. The model focuses on inactivated CD4+ T cells; it assumes that the T cell receptor (TCR) and its cofactors are active, and ignores the differences in glycolysis and lipid oxidation metabolism between effector and regulatory T cells. Furthermore, as the model is a minimal network, various components of the system were simplified, but previous simulations guarantee that the main dynamic regulatory motifs and feedback are considered (12). Given the available data, the model focuses on the observed behaviors in the VAT ignoring the contributions of other tissues. It also focuses on the first stage of hyperinsulinemia, ignoring long term effects, such as those presented under insulin resistance (9,34,36). The model does not include the dynamic interaction with adipocytes or macrophages, nor the effect of sexual hormones (15–17).

**Fig 1.**
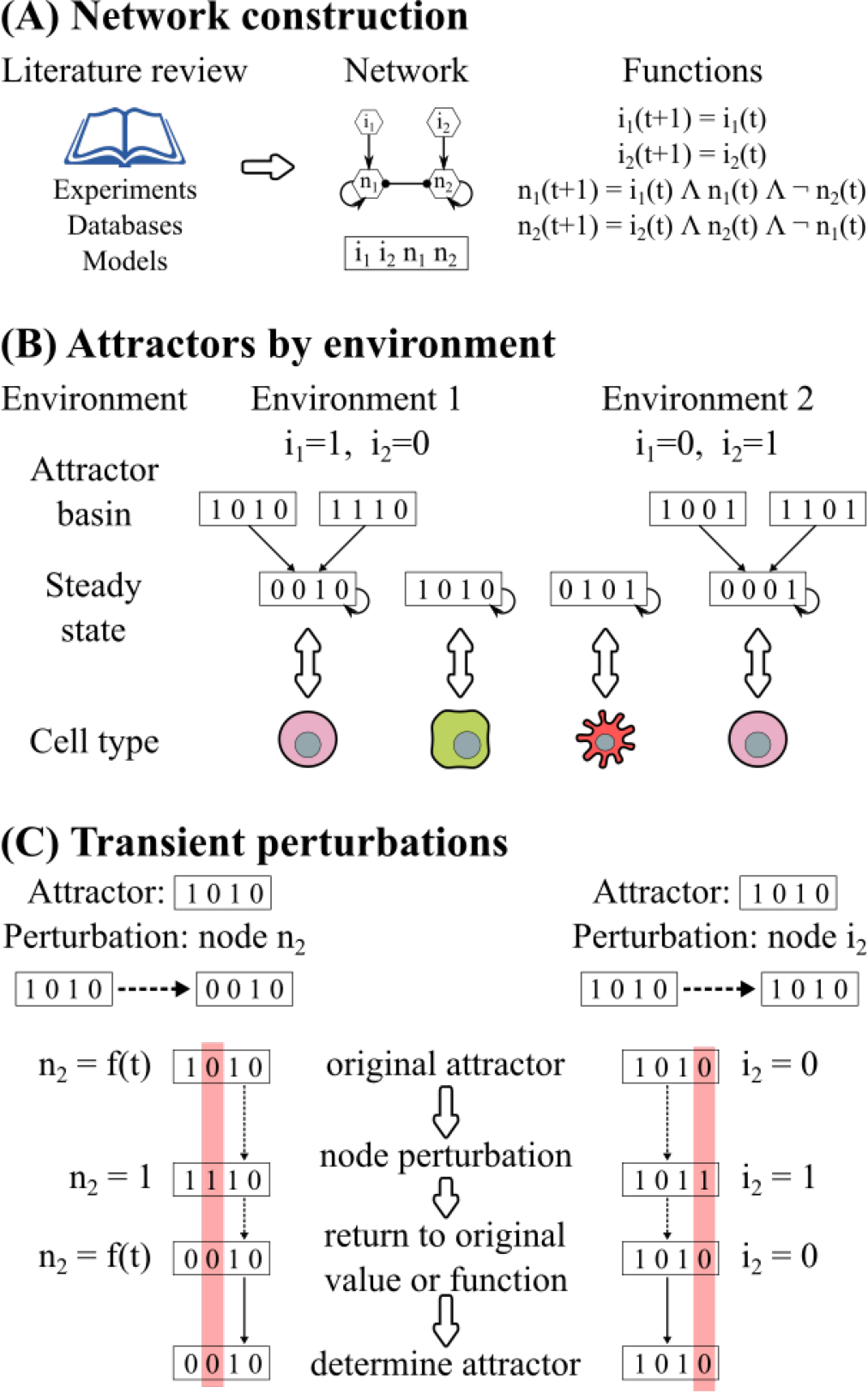
Experimental design of simulations. A) The network and regulatory functions were grounded on published experimental results. (B) The different inflammatory conditions were simulated by fixing the values of the input nodes of the network, that represent the extrinsic cytokines present in the microenvironment. For each simulated condition, the attractors and basins of attraction of the network were obtained. (C) The attractors of the network were perturbed by fixing the value of the target node for one time step and then returning the node to its original function or value; the system attractor was determined.

The nodes of the network correspond to transcription factors, signaling pathways and cytokines, while the edges correspond to the regulatory interactions between the nodes and are modeled as Boolean functions [Figure 1A; [Table S1]]. The resulting network contains 19 nodes and 54 interactions [Figure 2, BioModelsDatabase: MODEL1606020000]. The nodes include: transcription factors (Tbet, GATA3, RORγt, Foxp3), the effector and regulatory cytokines produced by the cell and their signaling pathways (intrinsic) (IFNγ, IL-2, IL-4, IL-21, TGFβ and IL-10), and the cytokines produced by the rest of the immune system (extrinsic) (IFNγe, IL-2e, IL-4e, IL-10e, IL-12e, IL-21e, IL-27e, and TGFβe). To simulate the effect of hyperinsulinemia we extended the previous network to add the regulation of IL-10 by insulin via the AKT pathway (14); and the STAT3-signaling cytokines: IL-10, IL-6, and IL-21 all use STAT3. We assumed that a different pathway mediates IL-10 signaling than IL-6/IL-21. As the model focuses on activated CD4+ T cells, we assume that the TCR signaling pathway is constitutively active and did not include explicitly this component in the network. The state of a node represents whether the biological component is active (1) or inactive (0). A node is active when it is capable of altering the regulation of other components of the immune system. For example, as CD4+ T cells require a basal level of insulin to survive, we considered this basal level to have a value of 0, while a higher insulin concentration, that is capable of affecting IL-10, is fixed to 1(14,37). In other words, hyperinsulinemia is simulated by setting the “insulin” node to 1.

**Fig 2.**
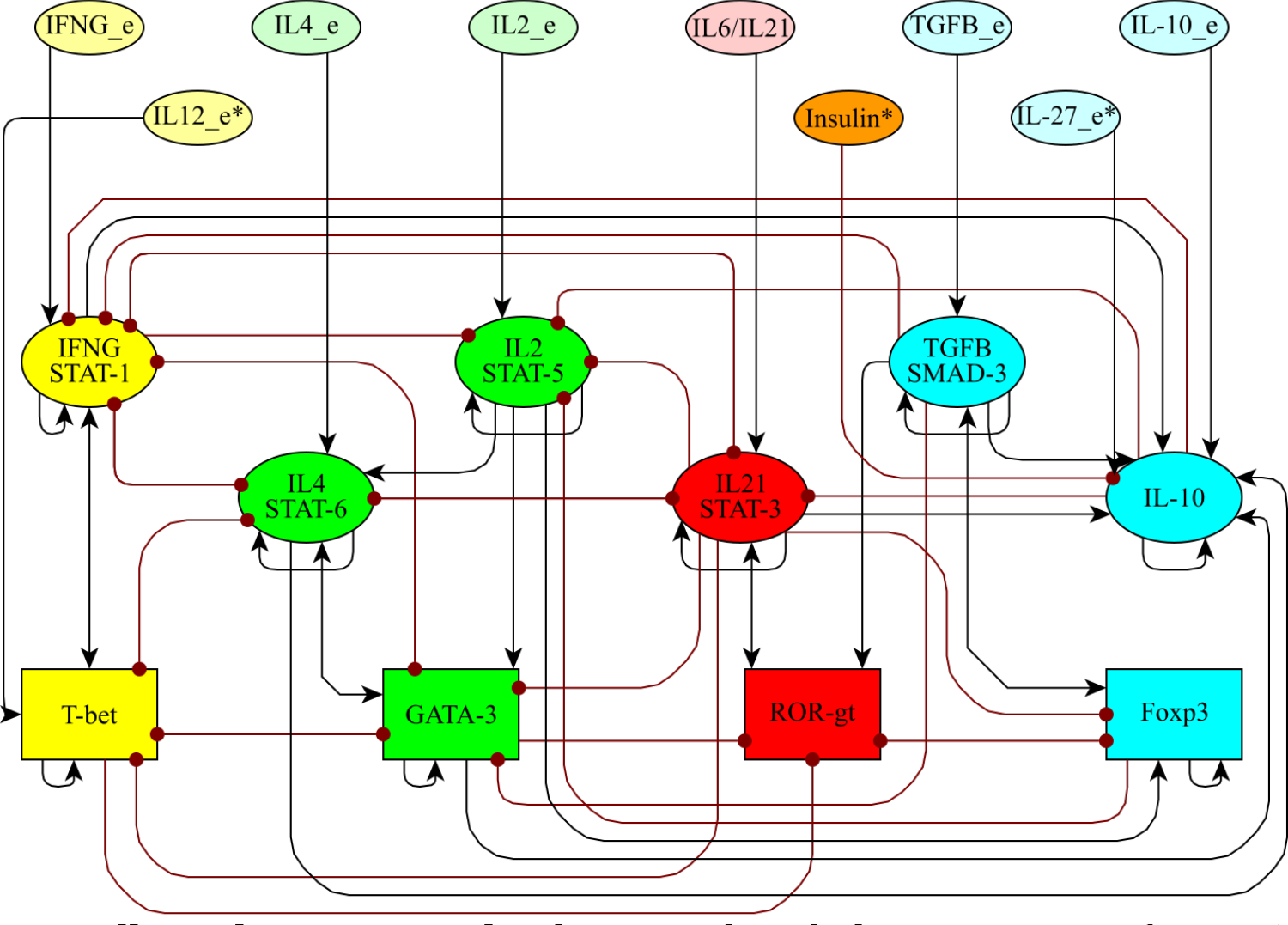
CD4+ T cell regulatory network. The network includes transcription factors (rectangles), intrinsic cytokines and their signaling pathways (ellipses), and extrinsic cytokines and insulin (ellipses).
Node colors correspond to cell types in which each molecule is generally expressed (state = 1): Th1 (yellow), Th2 (green), Th17 (red), iTreg (blue), and insulin (orange). Activations between elements are represented with black arrows, and inhibitions with red dotted arrows. An * is used to indicate the new nodes considered in this network model with respect to that in (12).

Cytokines can be produced by the cell (intrinsic) or by other cells of the immune system (extrinsic). Such extrinsic cytokines constitute the micro-environment and have an important role in CD4+ T cell differentiation and plasticity. Extrinsic cytokines were considered as inputs of the system [Figure 1B]. To study the effect of the micro-environment we focused on six biologically relevant environments: pro-Th0 or resting, pro-Th1, pro-Th2, pro-Th17, pro-iTreg, and pro-IL10 [Table 1].

**Table 1:**
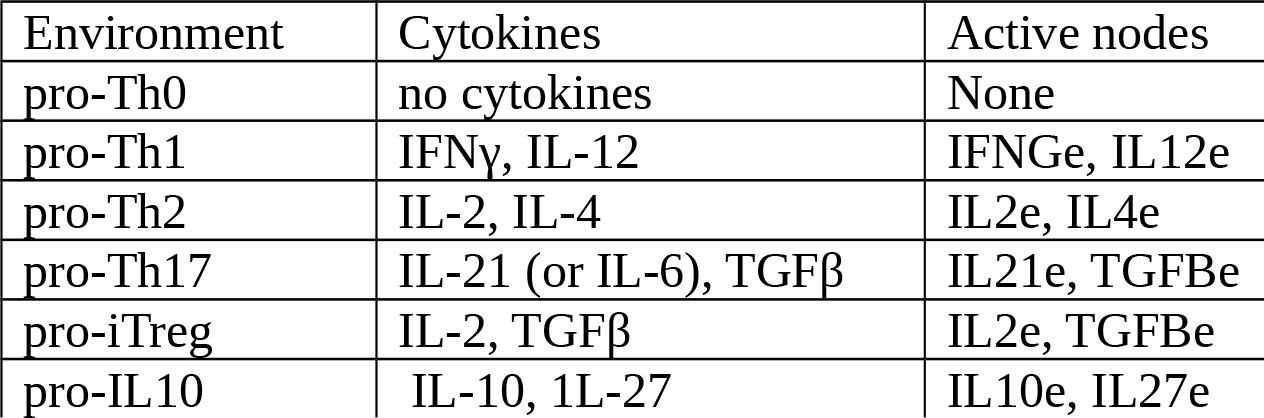
Environments of the CD4+ T cell regulatory network.

The stable states to which a regulatory network converge are called attractors, and can be interpreted as the expression profiles of the biological cell types (37,38) [Figure 1B]. We labeled each attractor according to the active transcription factors and intrinsic cytokines [Table S2]. Th0, resting T cells, were defined as expressing no transcription factors or regulatory cytokines. Th1 was defined as having Tbet and IFNγ active, Th2 as GATA3 and IL-4 active and GATA3+ (a Th2-like cell type) as GATA3+IL4-. Th17 cells are characterized by the expression of RORγt and STAT3 signaling mediated by IL-6 or IL-21, all of which require the presence of TGF-βe. The iTreg type has Foxp3 and TGFβ, IL-10 or both, all of which require the presence of IL-2e. T regulatory Foxp3-independent cells feature IL-10 (IL10+), TGF-β (TGFβ+) or both (IL10+TGFβ+), without expressing Foxp3. Th1 regulatory cells (Th1R) express a regulatory cytokine and T-bet [46]. Th2 regulatory cells (Th2rR) express a regulatory cytokine and GATA3. The attractors obtained by the CD4 + T cell network correspond to configurations that are characteristic of: Th0, Th1, Th1R, Th2, GATA3+, Th2R, Th17, iTreg, TGFβ+IL10+, TGFβ+ and IL10+ CD4+ T cells [Figure S1] {Zhu2010, MartinezSanchez2015}.

### Effect of insulin on CD4+ T cell differentiation

To simulate the effects of insulin, we obtained the attractors in the different micro-environments in the presence of basal levels (state of the “insulin” node to 0) or high levels (state of the “insulin” node to 1) of insulin [Figure 3]. To simulate the different environments we fixed the values of the input nodes according to each environment as listed in Table 1. Then, we determined and labeled the resulting attractors to obtain the predicted cell types under each environment and insulin condition [Figure 1B].

**Fig 3.**
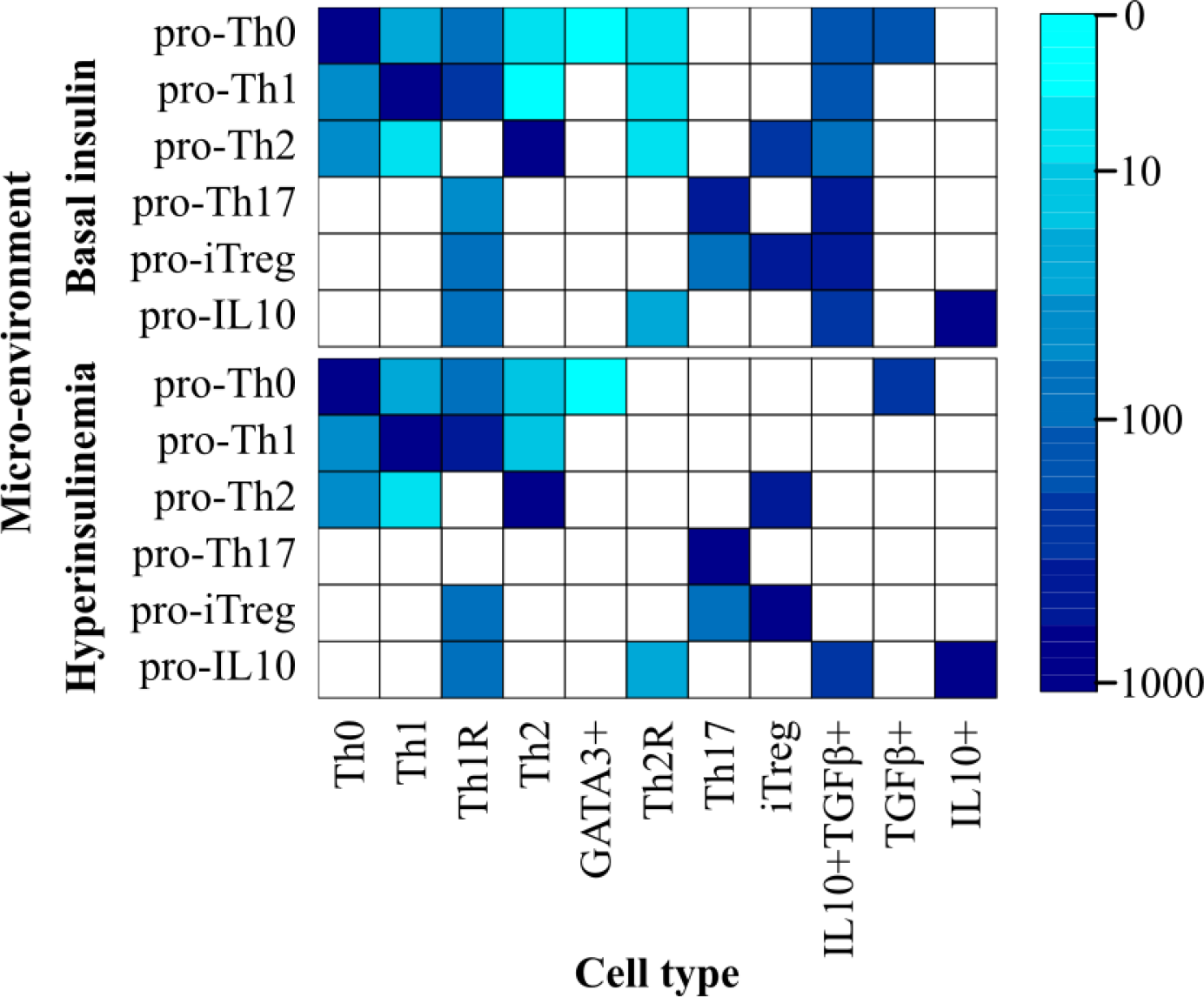
Effect of the micro-environment on CD4+ T cell differentiation. The values of the extrinsic signals of the TSRN were fixed according to different polarizing microenvironments. The color corresponds to the size of the basins of attraction on a logarithmic scale.

Our model shows that in effector polarizing environments with basal levels of insulin, like pro-Th1, pro-Th2 and pro-Th17, effector and regulatory cells coexist. In a pro-iTreg environment there is a coexistence of iTreg and Th17 cells. But in a pro-IL10 we see a strong polarization towards regulatory T cells and no effector CD4+ T cells. We observed that in the presence of high levels of insulin there is a marked decrease of the attractors that express IL-10 (Th1R, Th2R, and IL10+TGFβ+), and the remaining attractors tend to express TGFβ. There is an increase in the size of the basins of attraction of the Th17 and Th1 attractors. This is particularly notable in the pro-Th17 insulin environment, where the Th1R and IL10+TGFβ+ disappear, and the network converges to Th17. In the case of the Th1 attractor the increase in its basin size is smaller. Interestingly, this behavior corresponds to the observed increase in Th1 and Th17 and the decrease in Treg cells and IL-10 in obesity-associated chronic inflammation. The only exception to this pattern was observed under the pro-IL10 environment that remains unchanged by the level of insulin.

### Effect of insulin on CD4+ T cell plasticity

CD4+ T cells are plastic and dynamically change from one type to others, depending on the microenvironment and transient perturbations or initial conditions. This implies that these cells configurations and behavior can be altered dynamically. The multistable Boolean network model used here is a useful tool to study CD4+ T cell plasticity (12) as well. To explore this, we transiently perturbed the attractors for each microenvironment (Table 1) under the constitutive presence of basal (0) and high levels of insulin (1). For each attractor, we transiently perturbed each node for one time step. Then, we returned the node to its original value and used the original logical rules, to recover the resulting attractor [Figure 1C]. We established that a labeled attractor was robust to a perturbation if it returned to the same configuration after such transient perturbation, or to one that corresponds to the same cell type, after a perturbation. When the system transitioned to an attractor that corresponds to a different cell type, we considered the original attractor to be plastic under transient perturbations.

Our results suggest that the effect of insulin on the differentiation and plasticity of CD4+ T cells depends on the cytokines that are present in the microenvironment [Figure 4, File S2]. In each microenvironment, without insulin, most of the transitions lead the system to the favored cell type, which tends to be the most stable one, as expected. But in these cases, other cell types also coexist in the environment, especially regulatory cell types, even though the attractors that characterize them are less stable. Under high levels of insulin, that simulates an acute hyperinsulinemia condition, the CD 4+ T plasticity patterns are altered. In general, the activation of insulin: (1) causes the loss of the regulatory attractors, particularly those that express IL-10-, reduces cell stability, and the number of transitions towards the original cell type. This is particularly notable in the pro-Th17 environment, where Th17 is the only possible attractor. In the case of the pro-iTreg environment, there is a coexistence of iTreg with Th17. This is caused by the extrinsic TGFβ. The role of TGFβ is bivalent, as it can induce both regulation and inflammation through iTreg and Th17 cells. In the case of TGFβ, insulin shifts the equilibrium towards inflammatory cell types. The addition of insulin caused the loss of the IL10+TGFβ+ attractor, stabilized Th17, and reduced the stability of iTreg and Th1R. In the only case that this did not occur, was under the pro-IL10 environment, where a regulatory phenotype is attained independently of the insulin level, avoiding a pro-inflammatory condition even under the presence of hyperinsulinemia.

**Fig 4:**
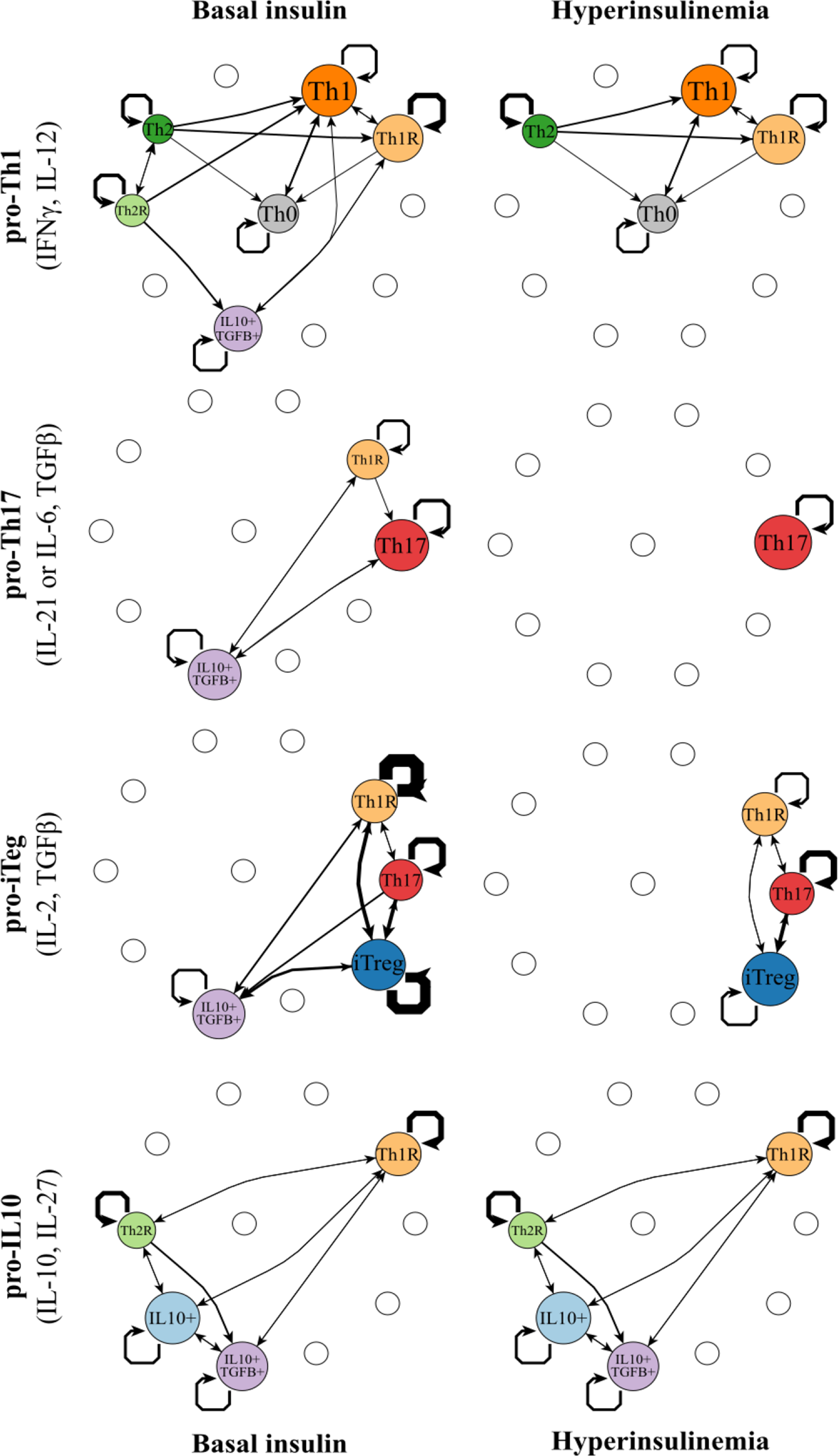
Cell fate map under different microenvironments. The values of the extrinsic signals of the CD4+ T regulatory network were fixed according to different polarizing micro environments as listed in Table 1, and the resulting attractors were transiently perturbed for one time step. Nodes correspond to cell types, node size is proportional to the number of configurations in a basin of attraction. Edges represent transitions from one cell type to another, their width represents the number of times the transition occurred, self-loops correspond to perturbations in which the network returned to the original cell type. The following micro-environments were studied:: (A) pro-Th1, (B) pro-Th1+ Insulin, (C) pro-Th17, (D) pro-Th17 + Insulin, (E) pro-iTreg, (F) pro-iTreg + Insulin, (G) pro-IL10 (H) pro-IL10.

### The role of IL10 on CD4+ T cell plasticity alterations under normal and hyperinsulinemic conditions

We assessed how many transitions among attractors were caused by transient perturbations of insulin and IL10 under normal and hyperinsulinemic conditions. On average perturbations of any node caused transitions to new cell types in 38% of the cases. But the number of transitions between cell types varied according to the node and the microenvironment. The transient increase of insulin caused transitions towards inflammatory or TGFB producing cell types under basal insulin level, while, as expected, under hyperinsulinemia the transient activation of insulin did not cause any further transitions. The attractors of the pro-Th17 environment with basal levels of insulin were very sensitive to perturbations in the insulin node, while the attractors found in the pro-iTreg and the pro-IL10 environment were robust to this perturbations. The transient activation of IL-10 caused transitions towards regulatory cell types. The attractors of the pro-Th1, pro-Th17 with basal levels of insulin and the pro-iTreg with high level of insulin environments were, on the other hand, very sensitive to transient perturbations of IL10. In environments with basal levels of insulin, the transient activation of IL-10 caused some transitions towards Th1 and Th2 cell types in pro-Th1 and pro-Th2 environments, respectively. In these cases, the transient activation of IL-10 was sufficient to destabilize the attractor but not to shift the network towards a regulatory cell type.

**Fig 5:**
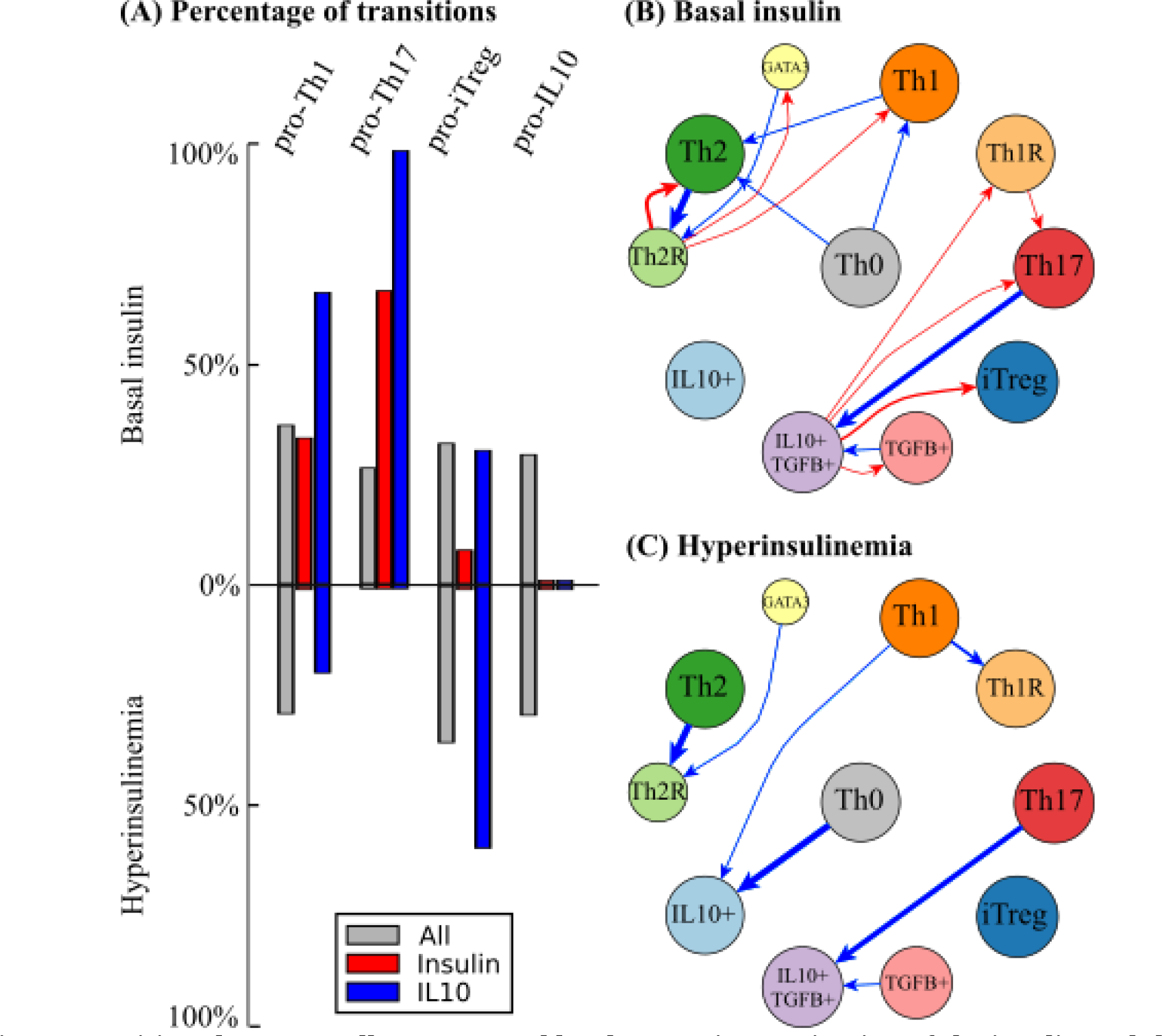
Transitions between cell types caused by the transient activation of the insulin and the IL-10 nodes. (A) Percentage of transitions between attractors in response to transient perturbations of: all nodes (grey); insulin (red) or IL-10 nodes (blue) under basal and high levels of insulin. (B, C) Transitions between cell types caused by the transient activation of: insulin (red) or IL-10 nodes (blue) under basal (B) and high levels (C) of insulin.

## Discussion

The theoretical simulation study presented here suggests that the impact of hyperinsulinemia on the inflammatory response (14), is mediated by the multistable dynamic GRN in (12). Overall, our simulation study provides a system-level platform to explain the relationship between hyperinsulinemia and altered proportions of T regulatory cells that have been observed in adipose tissue (3, 32–34, 36). It also highlights and provides a dynamic explanation to the different roles of TGFβ and IL-10 (25–28).

The model shows that in pro-Th1, pro-Th2, pro-Th17 and pro-iTreg microenvironments, effector and regulatory cells coexist. This pattern is observed in any disease, where it is common to observe cells from different subsets, even if a specific one is over-represented(39). Moreover, the simulations predict that different types of regulatory cells will predominate depending on the environment, being especially important to distinguish Foxp3+ and Fop3-regulatory T cells. Future experiments should consider that CD4+ T cells are highly heterogeneous, phenotypically plastic and sensitive to the microenvironment. A CD4+ T cell can express markers for more than one cell type at the same time, and its expression patterns can change over time, especially for regulatory T cells. It is necessary to measure the expression of Foxp3, IL-10, and TGFβ to systemically distinguish Treg (Foxp3+CD25high), Foxp3-IL10+, Foxp3-TGFβ+ and Th1R and Th2R hybrid cell types. Distinguishing these cell types will be necessary to understand the different roles that they play in obesity-associated chronic inflammation. Future assays should also consider multiple transcription factors and cytokines, carefully separate CD4+ T cell populations and compare their behaviors in different tissues.

Our simulation results recovered the altered CD4+ T cell populations that have been observed in murine models and humans during obesity-associated chronic inflammation (3,32–34,36). In the presence of hyperinsulinemia, increased proportions of Th1 and Th17 cells and decreased proportions of regulatory T cells are observed 3,33,34). Specifically, in a pro-Th17 environment, the presence of insulin predicts a complete shift towards Th17 cells. In contrast, in a pro-Th1 environment the Th1 attractor alteration is less dramatic than the alterations observed in vivo, probably because of the involvement of macrophages in the real condition, that are not considered in the simulation model of this study 3,33,34).

The model also provides an explanation to some paradoxical behaviors observed in CD4+ T regulatory cell populations during obesity-associated chronic inflammation. TGFβ can promote both inflammatory Th17 cells and regulatory Tregs, and transitions between both subsets have been observed 3,33,34,40). The model gives a mechanistic explanation to the fact that Th17 cells and iTregs are closely related and that Th17 cells can be observed sometimes during the iTreg response. TGFβ is necessary for the differentiation of both subsets, and transient signaling via the STAT3 pathway may be enough to shift some cells towards Th17, as the model shows. In obesity, Tregs expression profiles are similar to inflammatory T cells (32). Transfer and depletion of adipose Treg cells have been reported to both, improve or worsen insulin sensitivity, depending on the model and the population studied (33,35,36). Such apparently paradoxical behaviors can be explained by the relationship between TGFβ and IL-10 in the context of the dynamic regulatory network model used here. Under hyperinsulinemia, Th17 cells become more stable while IL10+ cells are lost. The remaining regulatory cells express TGFβ that is involved in Th17 differentiation, while insulin alters iTregs stability. In this way, the model predicts that hyperinsulinemic inflammatory environments, specially under pro-Th17 conditions, T regulatory cells are lost and the rest become unstable. In contrast, a pro-IL10 environment can induce regulatory T cells, regardless of the level of insulin in the environment. Nonetheless, while this pro-regulatory environment might decrease inflammation, it may have adverse effects as inflammation is relevant for the function of adipose tissue (42).

The model predicts that the transitions between cell types vary depending on the microenvironment and the perturbed node. Transient activation of insulin is sufficient to cause transitions towards inflammatory or TGFB+ cells, while transient activation of IL10 is sufficient to cause transitions towards regulatory cells. The stability of the different cell types will also vary depending on the microenvironment and the perturbation. We predict that the cells in a pro-Th17 environment are more sensitive to transient increases in insulin, while the cells in a pro-iTreg and pro-IL10 environments are more stable under this perturbation.

The model used here considers a minimum regulatory network underlying CD4+ T cell differentiation and plasticity under hyperinsulinemia, but it still lacks other cells and signals that are fundamental to fully understand obesity-associated chronic inflammation. For example, since the network used here is a minimal model, it ignores cytokines such as IL-1 and TNFα, the role of sexual hormones, and additional cell types such as adipocytes and macrophages, that play important roles during obesity-associated chronic inflammation (15–17). The model is restricted to assess the role of insulin on the differentiation dynamics of an activated CD4+ T cell in VAT, but still lacks the regulation of the TCR signaling pathway and the contrasting metabolism among effector, resting and regulatory conditions.

Furthermore, future efforts should consider continuous versions of the model to enable simulations of the strength and length of the signals in the dynamics of the immune system. Such simulations may be useful to assess different treatments of metabolic disorders and chronic inflammation, as well as the actual timing and progression of the obese inflammatory response. The model used here, still simplifies the microenvironment, that is more complex in vivo. For example, it is interesting to asses how the small initial signals that occur in response to nutrient overload, eventually give rise to significant alterations associated to obesity-associated chronic inflammation (15). Further studies of the effect of transient signals in a continuous version of the minimum and extended CD4+ T cell regulatory network, will likely yield important insights concerning such temporal patterns. Such system-level approach will be also useful for toxicological studies, and for providing predictions concerning the biological impact of drugs, assessing therapeutic targets or secondary effects.

## Materials and Methods

### Logical modeling formalism: Boolean networks

A Boolean network is composed of nodes that represent the system’s molecular components (i.e., cytokines, signaling pathways or transcription factors) and edges, that represent the interactions between nodes. The value of the nodes can be associated with a discrete variable denoting its current functional level of activity: if the node is functional its value is 1, and if it is not functional it is 0. The value of a node *x*_*i*_*(t+1)* depends on the value of its input nodes or regulators, this can be expressed with a Boolean function: 
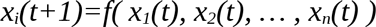

For the Th+insulin network, the Boolean functions were defined based on available T CD4+ differentiation models(10–12) and experimental data for the reported interactions among a network more than 90 nodes [Table S1]. The network was then simplified as (43) and GINSIM(44). The resulting network has 19 nodes and 54 interactions.

### Dynamic analysis

The state of the network *X* can be represented by a vector that specifies the value of all the nodes of the system. The state of the network will change over time depending on the Boolean functions associated with each node. When the values of a state vector *X* at *t*+1 are the same as those at time *t*+τ, the system has reached an attractor *X*\*:

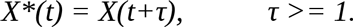

An attractor can be interpreted as a stable expression phenotype of a cell or cell type (45). All the states that lead to a solution *X*\* constitute the basin of attraction of such an attractor. We determined the stable states and basins of attraction of the network using GINSIM (44) and BoolNet(46).

### Labeling

Attractors were labeled depending on the expression of both the master transcription factors and cytokines. Labeling was automatized using BoolNetPerturb (47).

### Perturbations

To study the plasticity in response to perturbations we used BoolNetPerturb (47). First, we took all the attractors in each microenvironment, and systematically perturbed the value of the node for a time step, fixing the value of the target node during the corresponding time period. As the perturbation was transient, after a time step the node returns to its original function or-in the case of the inputs-to is original value. Finally, we reported the attractor that was reached after the perturbation. If the network returned to an attractor with the same label as the original attractor we said it was stable to that specific perturbation, if the network return to a different labeled attractor we said there had been a transition from one cell type to an other.

## Acknowledgments

This work is presented in partial fulfillment towards Mariana Martínez-Sanchez doctoral degree in the program “Doctorado en Ciencias Biomédicas, de la Universidad Nacional Autónoma de México”. We acknowledge the support Alejandro Frank and the “Centro de Ciencias de la Complejidad (C3), de la Universidad Nacional Autónoma de México”. We also acknowledge Diana Romo for her help with many logistical tasks and E. Roces de Álvarez-Buylla for her comments about this work. **Funding**: ERAB received funding from: CONACYT: 240180, 180380, 152649 and UNAM-DGAPA-PAPIIT: IN 203214, IN203814, IN211516. MI received funding from: PAPIIT IV100116. **Author contributions**: Conceived and designed the experiments: ERAB MEMS. Performed the experiments: MEMS. Analyzed the data: ERAB MEMS MH. Contributed reagents/materials/analysis tools: MEMS. Wrote the paper: MEMS ERAB MH. **Competing interests**: The authors declare that they have no competing interests. **Data and materials availability**: This model was deposited in BioModels and assigned the identifier M0DEL1606020000. The code and tutorials for the robustness and plasticity analysis are available at https://github.com/mar-esther23/boolnet-perturb.

## Supplementary Materials

**Fig S1:**
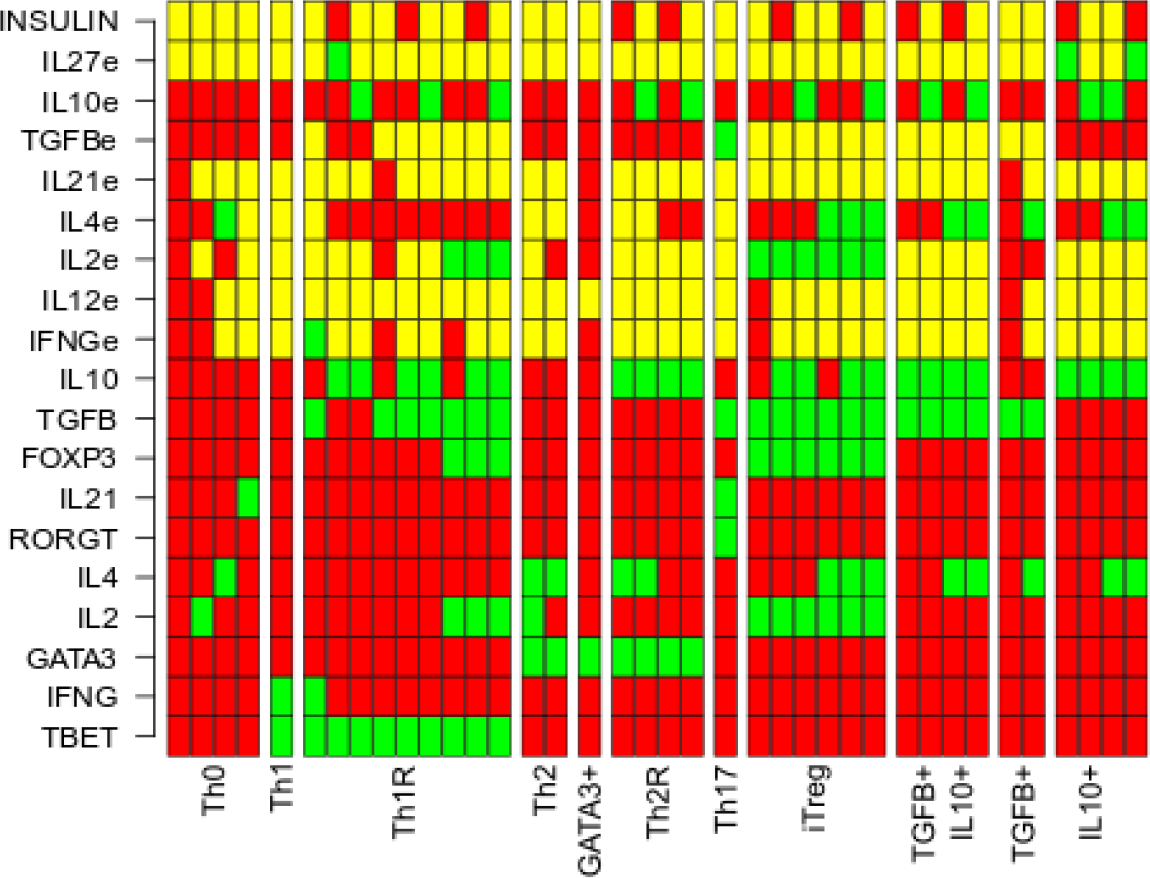
Attractors of the CD4+ T cell regulatory network. Each column corresponds to an attractor.Each node can be active (green) or inactive (red), extrinsic cytokines may be active or inactive (yellow). The following attractors were found in the network: Th0, Th1, Th1R, Th2, GATA3+, Th17, iTreg, TGFβ+IL10+, TGFβ+ and IL10+ regulatory cells. Attractors where labeled according to the active transcription factors and intrinsic cytokines.

**Table S1.**
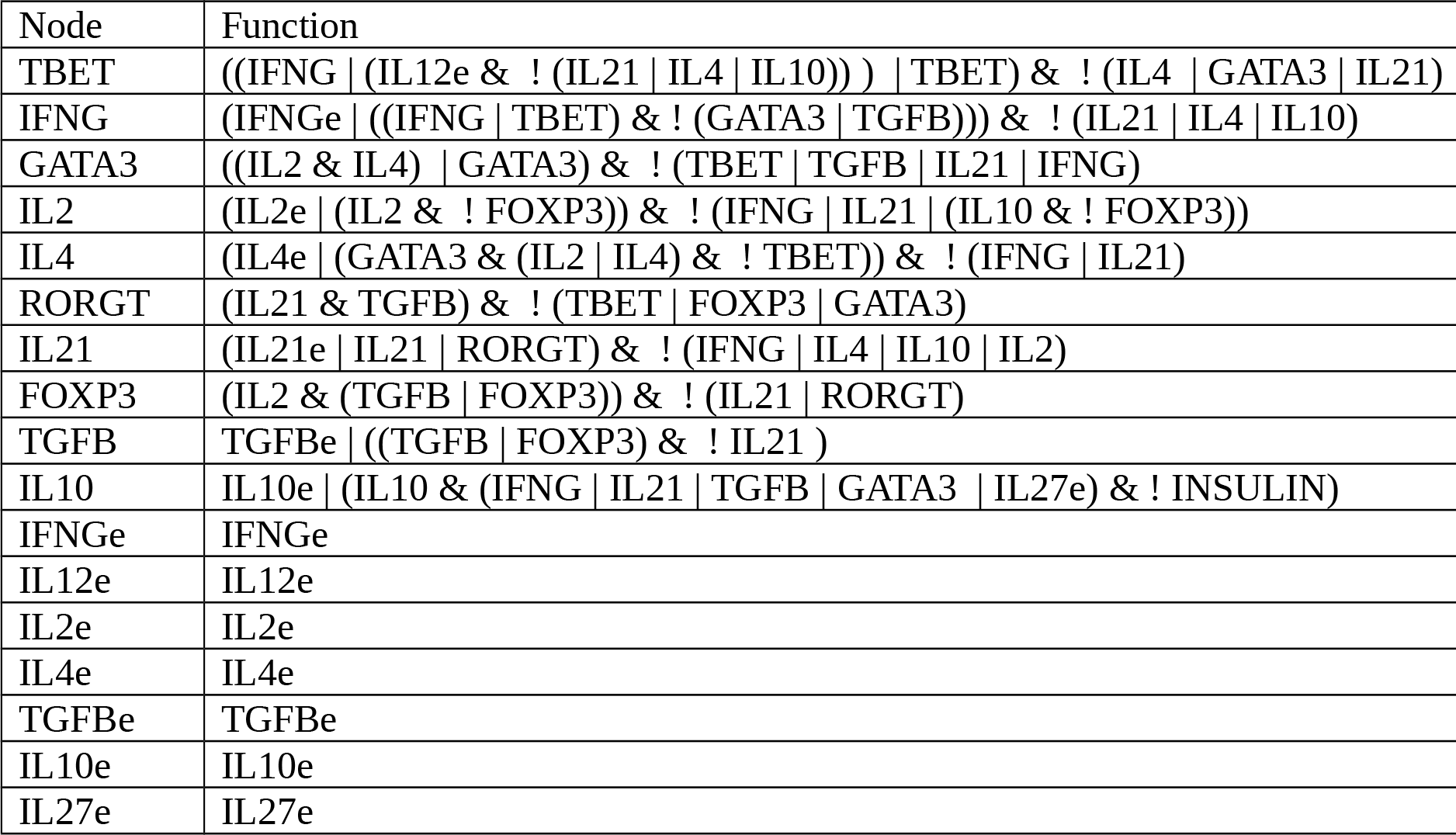
Rules of the CD4+ T cell regulatory network.

**Table S2.**
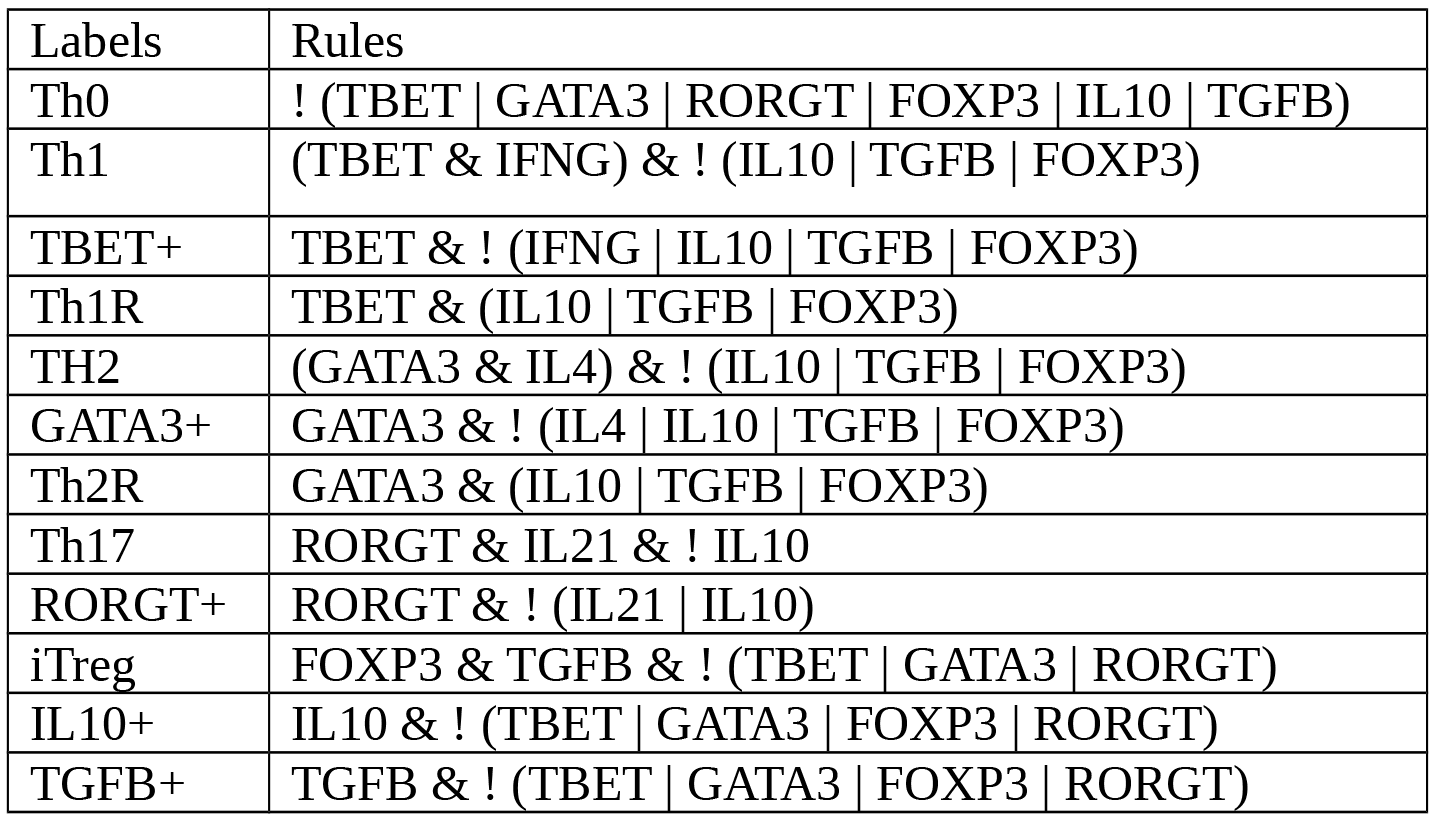
Labeling rules of the CD4+ T cell regulatory network.

## References

1. S. Winer and D. a Winer, “The adaptive immune system as a fundamental regulator of adipose tissue inflammation and insulin resistance,” Immunol. Cell Biol., vol. 90, no. 8, pp. 755–762, 2012.

2. J. M. Han and M. K. Levings, “Immune regulation in obesity-associated adipose inflammation.,” J. Immunol., vol. 191, no. 2, pp. 527–32, 2013.

3. V. A. Gerriets and N. J. MacIver, “Role of T cells in malnutrition and obesity.,” Front. Immunol., vol. 5, no. August, p. 379, 2014.

4. M. Hiriart, M. Velasco, C. Larque, and C. M. Diaz-Garcia, “Metabolic syndrome and ionic channels in pancreatic beta cells,” Vitam. Horm., vol. 95, pp. 87–114, 2014.

5. E. Balleza, E. R. Alvarez-Buylla, A. Chaos, S. Kauffman, I. Shmulevich, and M. Aldana, “Critical dynamics in genetic regulatory networks: examples from four kingdoms.,” PLoS One, vol. 3, no. 6, p. e2456, Jan. 2008.

6. S. Huang, “Systems biology of stem cells: three useful perspectives to help overcome the paradigm of linear pathways.,” Philos. Trans. R. Soc. Lond. B. Biol. Sci., vol. 366, no. 1575, pp. 2247–59, Aug. 2011.

7. P. Creixell, J. Reimand, S. Haider, G. Wu, T. Shibata, M. Vazquez, V. Mustonen, A. Gonzalez-Perez, J. Pearson, C. Sander, B. J. Raphael, D. S. Marks, B. F. F. Ouellette, A. Valencia, G. D. Bader, P. C. Boutros, J. M. Stuart, R. Linding, N. Lopez-Bigas, and L. D. Stein, “Pathway and network analysis of cancer genomes,” Nat. Methods, vol. 12, no. 7, pp. 615–621, 2015.

8. S. M. Hill, L. M. Heiser, T. Cokelaer, M. Unger, N. K. Nesser, D. E. Carlin, Y. Zhang, A. Sokolov, E. O. Paull, C. K. Wong, K. Graim, A. Bivol, H. Wang, F. Zhu, B. Afsari, L. V Danilova, A. V Favorov, W. S. Lee, D. Taylor, C. W. Hu, B. L. Long, D. P. Noren, A. J. Bisberg, B. Afsari, R. Al-Ouran, B. Anton, T. Arodz, O. A. Sichani, N. Bagheri, N. Berlow, A. J. Bisberg, A. Bivol, A. Bohler, J. Bonet, R. Bonneau, G. Budak, R. Bunescu, M. Caglar, B. Cai, C. Cai, D. E. Carlin, A. Carlon, L. Chen, M. F. Ciaccio, T. Cokelaer, G. Cooper, C. J. Creighton, S.-M.-H. Daneshmand, A. de la Fuente, B. Di Camillo, L. V Danilova, J. Dutta-Moscato, K. Emmett, C. Evelo, M.-K. H. Fassia, A. V Favorov, E. J. Fertig, J. D. Finkle, F. Finotello, S. Friend, X. Gao, J. Gao, J. Garcia-Garcia, S. Ghosh, A. Giaretta, K. Graim, J. W. Gray, R. GroEeholz, Y. Guan, J. Guinney, C. Hafemeister, O. Hahn, S. Haider, T. Hase, L. M. Heiser, S. M. Hill, J. Hodgson, B. Hoff, C. H. Hsu, C. W. Hu, Y. Hu, X. Huang, M. Jalili, X. Jiang, T. Kacprowski, L. Kaderali, M. Kang, V. Kannan, M. Kellen, K. Kikuchi, D.-C. Kim, H. Kitano, B. Knapp, G. Komatsoulis, H. Koeppl, A. Krämer, M. B. Kursa, M. Kutmon, W. S. Lee, Y. Li, X. Liang, Z. Liu, Y. Liu, B. L. Long, S. Lu, X. Lu, M. Manfrini, M. R. A. Matos, D. Meerzaman, G. B. Mills, W. Min, S. Mukherjee, C. L. Müller, R. E. Neapolitan, N. K. Nesser, D. P. Noren, T. Norman, B. Oliva, S. O. Opiyo, R. Pal, A. Palinkas, E. O. Paull, J. Planas-Iglesias, D. Poglayen, A. A. Qutub, J. Saez-Rodriguez, F. Sambo, T. Sanavia, A. Sharifi-Zarchi, J. Slawek, A. Sokolov, M. Song, P. T. Spellman, A. Streck, G. Stolovitzky, S. Strunz, J. M. Stuart, D. Taylor, J. Tegnér, K. Thobe, G. M. Toffolo, E. Trifoglio, M. Unger, Q. Wan, H. Wang, L. Welch, C. K. Wong, J. J. Wu, A. Y. Xue, R. Yamanaka, C. Yan, S. Zairis, M. Zengerling, H. Zenil, S. Zhang, Y. Zhang, F. Zhu, Z. Zi, G. B. Mills, J. W. Gray, M. Kellen, T. Norman, S. Friend, A. A. Qutub, E. J. Fertig, Y. Guan, M. Song, J. M. Stuart, P. T. Spellman, H. Koeppl, G. Stolovitzky, J. Saez-Rodriguez, and S. Mukherjee, “Inferring causal molecular networks: empirical assessment through a community-based effort,” Nat. Methods, no. February 2015, 2016.

9. J. M. Olefsky and C. K. Glass, “Macrophages, inflammation, and insulin resistance.,” Annu. Rev. Physiol., vol. 72, pp. 219–46, Jan. 2010.

10. A. Carbo, R. Hontecillas, B. Kronsteiner, M. Viladomiu, M. Pedragosa, P. Lu, C. W. Philipson, S. Hoops, M. Marathe, S. Eubank, K. Bisset, K. Wendelsdorf, A. Jarrah, Y. Mei, and J. Bassaganya-Riera, “Systems modeling of molecular mechanisms controlling cytokine-driven CD4+ T cell differentiation and phenotype plasticity.,” PLoS Comput. Biol., vol. 9, no. 4, p. e1003027, Apr. 2013.

11. W. Abou-Jaoudé, P. T. Monteiro, A. Naldi, M. Grandclaudon, V. Soumelis, C. Chaouiya, and D. Thieffry, “Model checking to assess T-helper cell plasticity.,” Front. Bioeng. Biotechnol., vol. 2, p. 86, Jan. 2014.

12. M.E. Martinez-Sanchez, L. Mendoza, C. Villarreal, and E.R. Alvarez-Buylla, “A Minimal Regulatory Network of Extrinsic and Intrinsic Factors Recovers Observed Patterns of CD4+ T Cell Differentiation and Plasticity.,” PLoS Comput. Biol., vol. 11, no. 6, p. e1004324, Jul. 2015.

13. N. J. MacIver, R. D. Michalek, and J. C. Rathmell, “Metabolic Regulation of T Lymphocytes.,” Annu. Rev. Immunol., vol. 31, no. December 2012, pp. 259–283, Jan. 2013.

14. J. M. Han, S. J. Patterson, M. Speck, J. a. Ehses, and M. K. Levings, “Insulin inhibits IL-10-mediated regulatory T cell function: implications for obesity.,” J. Immunol., vol. 192, no. 2, pp. 623–9, Jan. 2014.

15. M. F. Gregor and G. S. Hotamisligil, “Inflammatory mechanisms in obesity.,” Annu. Rev. Immunol., vol. 29, pp. 415–45, Apr. 2011.

16. M. Hiriart, C. L., Myrian Velasco, Carlos Manlio Diaz-Garcia, A., Carmen Sanchez-Soto, J. P. C. Albarado-Ibaez, and G. nvez-Maldonado, Alicia Toledo and Neivys Garci a-DelgadoKanakis, “Pancreatic Beta Cells in Metabolic Syndrome,” in Islets of Langerhans SE–50, 2014, pp. 1–29.

17. J. Hirosumi, G. Tuncman, L. Chang, C. Z. Gorgun, K. T. Uysal, K. Maeda, M. Karin, and G. S. Hotamisligil, “A central role for JNK in obesity and insulin resistance.,” Nature, vol. 420, no. 6913, pp. 333–6, Dec. 2002.

18. C. T. Kelly, J. Mansoor, G. L. Dohm, W. H. H. Chapman, J. R. Pender, and W. J. Pories, “Hyperinsulinemic syndrome: The metabolic syndrome is broader than you think,” Surg. (United States), vol. 156, no. 2, pp. 405–411, 2014.

19. J. Zhu, H. Yamane, W. E. W. Paul, and Z. J. Y. H. P. We, “Differentiation of effector CD4 T cell populations,” Annu. Rev. Immunol., vol. 28, no. 1, pp. 445–89, Jan. 2010.

20. T. Hong, J. Xing, L. Li, and J. J. Tyson, “A simple theoretical framework for understanding heterogeneous differentiation of CD4+ T cells.,” BMC Syst. Biol., vol. 6, no. 1, p. 66, Jan. 2012.

21. S. Huang, “Hybrid T-Helper Cells: Stabilizing the Moderate Center in a Polarized System,” PLoS Biol., vol. 11, no. 8, pp. 1–5, 2013.

22. K. K. M. K. M. Murphy, B. Stockinger, and A. Manuscript, “Effector T cell plasticity: flexibility in the face of changing circumstances,” Nat. Immunol., vol. 11, no. 8, pp. 674–680, Aug. 2010.

23. S. Sakaguchi, D. a a Vignali, A. Y. Rudensky, R. E. Niec, and H. Waldmann, “The plasticity and stability of regulatory T cells.,” Nature reviews. Immunology, vol. 13, no. 6. pp. 461–7, 2013.

24. P. We, “Fundamental Immunology 6th ed.” Lippincott Williams & Wilkins, Reading, Massachusetts, 2008.

25. F. Fantini MC Becker C, “Cutting edge: TGF-β induces a regulatory phenotype in CD4+CD25-T cells through Foxp3 induction and down-regulation of Smad7,” J Immunol, vol. 172, pp. 5149–5153, 2004.

26. B. Veldhoen M Hocking RJ, M. Veldhoen, R. J. Hocking, C. J. Atkins, R. M. Locksley, and B. Stockinger, “TGFβ in the context of an inflammatory cytokine milieu supports de novo differentiation of IL-17-producing T cells,” Immunity, vol. 24, no. 2, pp. 179–189, Feb. 2006.

27. M. a Travis and D. Sheppard, “TGF-β activation and function in immunity.,” Annu. Rev. Immunol., vol. 32, no. November 2013, pp. 51–82, 2014.

28. A. Howes, P. Stimpson, P. Redford, L. Gabrysova, and A. O’Garra, “Interleukin-10: Cytokines in Anti-inflammation and Tolerance,” in Cytokine Frontiers: Regulation of Immune Responses in Health and Disease, vol. 6, 2014, pp. 327-352.

29. M. a Koch, G. Tucker-Heard, N. R. Perdue, J. R. Killebrew, K. B. Urdahl, D. J. Campbell, “The transcription factor T-bet controls regulatory T cell homeostasis and function during type 1 inflammation.,” Nat. Immunol., vol. 10, no. 6, pp. 595–602, Jun. 2009.

30. M. J. Barnes and F. Powrie, “Hybrid T reg cells: steel frames and plastic exteriors,” Immunity, vol. 10, no. 6, pp. 11–12, 2009.

31. Y. Wang, M. A. Su, and Y. Y. Wan, “An essential role of the transcription factor GATA-3 for the function of regulatory T cells.,” Immunity, vol. 35, no. 3, pp. 337–348, Sep. 2011.

32. M. Feuerer, L. Herrero, D. Cipolletta, A. Naaz, J. Wong, J. Lee, A. Goldfine, C. Benoist, S. Shoelson, and D. Mathis, “Fat Treg cells: a liaison between the immune and metabolic systems,” Nat. Med., vol. 15, no. 8, pp. 930–939, 2009.

33. O. Osborn and J. M. Olefsky, “The cellular and signaling networks linking the immune system and metabolism in disease,” Nat. Med., vol. 18, no. 3, pp. 363–374, 2012.

34. T. McLaughlin, L. F. Liu, C. Lamendola, L. Shen, J. Morton, H. Rivas, D. Winer, L. Tolentino, O. Choi, H. Zhang, M. H. Y. Chng, and E. Engleman, “T-cell profile in adipose tissue is associated with insulin resistance and systemic inflammation in humans,” Arterioscler. Thromb. Vasc. Biol., vol. 34, no. 12, pp. 2632–2636, 2014.

35. M. Feuerer, L. Herrero, D. Cipolletta, A. Naaz, J. Wong, A. Nayer, J. Lee, A. B. Goldfine, C. Benoist, S. Shoelson, and D. Mathis, “Lean, but not obese, fat is enriched for a unique population of regulatory T cells that affect metabolic parameters.,” Nat. Med., vol. 15, no. 8, pp. 930–9, Aug. 2009.

36. S. P. Bapat, J. Myoung Suh, S. Fang, S. Liu, Y. Zhang, A. Cheng, C. Zhou, Y. Liang, M. LeBlanc, C. Liddle, A. R. Atkins, R. T. Yu, M. Downes, R. M. Evans, and Y. Zheng, “Depletion of fat-resident Treg cells prevents age-associated insulin resistance,” Nature, 2015.

37. E. Azpeitia, M. Bemtez, P. Padilla-Longoria, C. Espinosa-Soto, and E.R. Alvarez-Buylla, “Dynamic network-based epistasis analysis: boolean examples.,” Front. Plant Sci., vol. 2, p. 92, Jan. 2011.

38. R. Albert and J. Thakar, “Boolean modeling: a logic-based dynamic approach for understanding signaling and regulatory networks and for making useful predictions,” Wiley Interdiscip. Rev. Syst. Biol. Med., vol. 6, no. 5, pp. 353–369, Sep. 2014.

39. M. DuPage and J. A. Bluestone, “Harnessing the plasticity of CD4+ T cells to treat immune-mediated disease,” Nat. Rev. Immunol., vol. 16, no. 3, pp. 149–163, Feb. 2016.

40. Y. K. Lee, R. Mukasa, R. D. Hatton, and C. T. Weaver, “Developmental plasticity of Th17 and Treg cells.,” Curr. Opin. Immunol., vol. 21, no. 3, pp. 274–80, Jun. 2009.

41. M. Kleinewietfeld and D. A. Hafler, “The plasticity of human Treg and Th17 cells and its role in autoimmunity.,” Semin. Immunol., vol. 25, no. 4, pp. 305–12, Nov. 2013.

42. I. Wernstedt Asterholm, C. Tao, T. S. Morley, Q. A. Wang, F. Delgado-Lopez, Z. V Wang, and P. E. Scherer, “Adipocyte inflammation is essential for healthy adipose tissue expansion and remodeling.,” Cell Metab., vol. 20, no. 1, pp. 103–18, Jul. 2014.

43. J. Davila-Velderrain, C. Villarreal, and E.R. Alvarez-Buylla, “Reshaping the epigenetic landscape during early flower development: induction of attractor transitions by relative differences in gene decay rates.,” BMC Syst. Biol., vol. 9, no. 1, p. 20, 2015.

44. A. Naldi, D. Berenguier, A. Fauré, F. Lopez, D. Thieffry, and C. Chaouiya, “Logical modelling of regulatory networks with GINsim 2.3.,” Biosystems., vol. 97, no. 2, pp. 134–9, Aug. 2009.

45. S. A. Kauffman, “Metabolic stability and epigenesis in randomly constructed genetic nets,” J. Theor. Biol., vol. 22, no. 3, pp. 437–467, Mar. 1969.

46. C. Müssel, M. Hopfensitz, and H. A. Kestler, “BoolNet-an R package for generation, reconstruction and analysis of Boolean networks,” Bioinformatics, vol. 26, no. 10, pp. 1378–1380, 2010.

47. M. E. Martinez-Sanchez. BoolNet Perturb, GitHub repository:https://github.com/mar-esther23/boolnet-perturb. 2015

48. Chelliah V et al. BioModels: ten-year anniversary. Nucl. Acids Res. 2015, 43(Database issue):D542–8

